# Low rate of follow-up colonoscopy after positive results of colorectal cancer screening in a Chinese urban core district

**DOI:** 10.1101/734152

**Authors:** Yawen Guo, Qingwu Jiang, Tetsuya Tanimoto, Masahiro Kami, Peng Peng, Yongpin He, Xiaoming Yang, Xin Zhang, Wenjun Gao, Yanming Wang, Xiaoting Chu, Yunhui Wang, Shigeaki Kato, Xiaocao Ding

## Abstract

Colorectal cancer (CRC) is one of the most common cancers in China. In 2003, a systematic CRC registry that enables the determination of CRC incidence and mortality and a CRC screening project were introduced in the Jing’an district of Shanghai by the municipal government. This study assessed the registry data to determine the status of CRC and CRC screening rates in the core district of an urban area of China. Data were retrieved from the Official registry information systems of Jing’an district Shanghai Cance. The incidence and mortality of CRC, as well as population-based CRC screening rates, were analysed. Individuals who screened positive for CRC based upon results of a high-risk factor questionnaire (HRFQ) and a faecal immunochemical test (FIT) were selected for follow-up colonoscopy (CSPY). From the registry data collected between 2003 and 2013, the standardized incidence rate was 26.44/10^5^, with a significant gender difference. The CRC standardized mortality rate was 10.08/10^5^. In 2013, 17,866 individuals (21.6%) enrolled for CRC screening among the 82,602 candidate residents. The positive screening rate was 16.28% (2909/17866). Among the 2909 positives, 508 (17.5%) underwent follow-up CSPY. In 41.3% of these individuals (210/508), abnormal lesions were detected. Of these, 8 (1.57%) lesions were diagnosed as CRC, and 142 (28.0%) were identified as precancerous lesions. During the assessment period, both the incidence and mortality of CRC in the Jing’an district were determined in the area of high CRC prevalence in Chin. Nevertheless, the rate of participation in CRC screening was low (21.6%), and the rate of participation in follow-up CSPY for individuals who screened positive was only 17.5%. Improved participation in CRC screening and follow-up CSPY is expected to lower the incidence and mortality of CRC significantly in the rural areas of China. (288)

## Introduction

Colorectal cancer (CRC) is one of the most commonly diagnosed cancers in males and in females worldwide.[1-3] CRC has a high prevalence, and it has become widely viewed as having a long latency period, providing ample time for both early detection and prevention. Indeed, if diagnosed at the earliest stage, CRC can be treated successfully, and most patients will survive for more than five years.

Research has shown that CRC screening reduces the incidence and mortality of CRC. A meta-analysis of trials demonstrated that CRC mortality could be reduced by 16% for those invited to screening.[4] Based on the effectiveness of screening, a guideline for screening and surveillance for the early detection of CRC and adenomatous polyps was published jointly by the American Cancer Society, the US Multi-society Task Force on CRC, and the American College of Radiology in 2008.[5] Similar systems and guidelines for CRC screening have been established in other Western and Asian countries as well.[6-12] In China, several large-scale population screenings using the Sequential Fecal Occult Blood Screening Program were performed by the municipals, and the public data reported in the several Chinese medical journals showed high rates of CRC and adenomatous polyps that were similar to those reported in other countries.[13-16] Furthermore, to facilitate CRC screening, a high-risk factor questionnaire (HRFQ) was generated and applied for the CRC screening from 1992. However, as there is huge diversity in the urban as well as the rural areas in China, mainly derived from the availability of medical resources and mobility of the residents, little information was available for accurate assessment of the CRC incidence and mortality of permanent residents.

In the urban areas of Shanghai, CRC was ranked higher in incidence (second) and mortality (fourth) among all malignant tumours from the public data source released from the municipals. The Jing’an district is small (0.12% of the total Shanghai area), but it is located in the core and most fruitful centre of Shanghai in terms of economical and medical resources, with a higher life expectancy than the entire Shanghai area in 2015. Owing to this situation, the residents of this district tend to live there permanently. Therefore, this district appears appropriate for assessment of CRC incidence and mortality, as well as the rate of participation in CRC screening, as a representative urban area in China.

To accurately assess CRC incidence and mortality and the rate of participation in CRC screening in urban areas, the present study was conducted by retrieving the related informations of the residents of the Jing’an district from the Registry Information Systems in Shanghai for analyses. The results showed a high CRC incidence and mortality during the assessment period, together with a low participation rate in CRC screening (21.6%). Notably, only 17.5% (508) of the 2,909 individuals who screened positive for CRC were found to take follow-up colonoscopy (CSPY) in 2013.

## Materials and methods

### Data source for CRC incidence and mortality assessment

In Shanghai, cause of death monitoring began in the 1950s. The Shanghai All Causes of Death Registration Reporting System was established in every district of Shanghai in 1973. The Cancer Prevention and Control Office was established in 1986. In 2001, the Shanghai Municipal Health and Family Planning Commission launched the Shanghai Malignant Tumor Report Methods. Since then, hospital tumour registration and reporting has been standardized, and the Shanghai Cancer Registry Information System has been used. We retrieved the related informations from the Jing’an district CDC (Shanghai Jingan Center for Disease Control and Prevention), through the Shanghai Cancer Registry Information System and the Shanghai All Causes of Death Registration Reporting System organized by the Shanghai Government and Jingan municipal office.

The CRC screening was conducted by six sites of the community health service center in this district under supervise by the health bureau of the Shanghai Government and Jingan municipal office. Colonoscopy and the follow-up diagnosis for further treatment under supervise by the health bureau were performed in the four designated hospitals (Jing’an District Central hospital, Huashan hospital, Huadong hospital and Post & Telecommunications Hospital of Shanghai.).

### Incidence and mortality rate and standardization

Data for the registered population was obtained from the Jing’an District Public Security Bureau to calculate the crude incidence (mortality) rate. We used the 1966 world standard population composition [17] to calculate the world population-adjusted incidence (mortality), which was referred to as the standardized rate.

### Data source for screening

The Shanghai CRC Screening Information System has been used since 2013, when population-based CRC screening was introduced in Shanghai. We obtained the data from the Shanghai CRC Screening Information System of the Jing’an district. The project population consisted of the resident population from 50 to 74 years of age, and 82,602 persons were offered screening based on China’s sixth census of the resident population in the Jing’an district.

### Evaluation

Based on the Shanghai Community Residents CRC Screening Program, we adopted a two-step protocol for population-based CRC screening: (1) an initial high-risk factor questionnaire (HRFQ) and faecal immunochemical test (FIT) followed by (2) a full CSPY for those cases identified as positive in the initial screening. When participants with either a positive HRFQ or one of two FITs proved positive, they were referred for CSPY.

### HRFQ

According to the “CRC Risk Assessment Form for Shanghai Community Residents”, we evaluated the CRC risk of the target population and screened out high-risk groups by: (1) first-degree relative(s) with CRC; (2) personal history of cancer; (3) personal history of intestinal polyps; (4) two or more of the following symptoms/histories: (4a) chronic diarrhoea (diarrhoea lasting for more than 3 months in the last two years, and each episode lasting for more than 1 week); (4b) chronic constipation (over the past two years, constipation lasting for more than two months every year); (4c) mucous and bloody stool; (4d) major trauma or painful event (e.g. divorce, death of relatives); (4e) history of chronic appendicitis or appendectomy; (4f) history of chronic cholecystitis or cholecystectomy. If an individual was positive for any one of the 4 items above, the result was considered positive on the HRFQ.

### Examinations

The FIT was performed 2 times per person at a one-week interval. Participants were informed of the significance of the positive results and given a referral for CSPY to be carried out at a designated hospital. These designated hospitals were asked to input CSPY information in a timely manner.

### Quality control

A random sample of 2% each day was checked, and the coincidence rate was not less than 90%. After the original questionnaire information was entered, the coincidence rate of the information system and the questionnaire should be 100%.

### Statistical analyses

Categorical data were analysed using the Chi-square test, and a difference of *P*<0.05 was considered statistically significant. Using SAS 9.1.3, we compared detection rates with Fisher’s Exact Test (two-sided) on both sides of the Exact *P*-value, recording the *P*-value of the Chi-square at the same time. Chi-square was analysed by age group, using hierarchical analysis.

## Results

### Incidence and mortality of CRC in the Jing’an district from 2003 to 2013

The crude and standardized incidence rates of CRC between 2003 and 2013 were 66.18/10^5^ and 26.44/10^5^ (Table 1).

**Table 1.** CRC incidence rate by year and sex (2003-2013) (/100 thousand)

|  | male |  |  |  |  |  | female |  |  |  |  |  | total |  |
| --- | --- | --- | --- | --- | --- | --- | --- | --- | --- | --- | --- | --- | --- | --- |
|  | Colon cancer |  | Rectal cancer |  | total |  | Colon cancer |  | Rectal cancer |  | total |  |  |  |
|  | crude | standard | crude | standard | crude | standard | crude | standard | crude | standard | crude | standard | crude | standard |
| 2003 | 32.35 | 14.42 | 21.15 | 10.45 | 53.51 | 24.88 | 35.58 | 13.49 | 15.68 | 7.28 | 51.26 | 20.77 | 52.37 | 23.04 |
| 2004 | 39.01 | 19.27 | 30.05 | 12.59 | 69.06 | 31.86 | 36.39 | 13.23 | 21.59 | 9.40 | 57.97 | 22.63 | 63.42 | 26.82 |
| 2005 | 37.80 | 17.47 | 26.72 | 14.47 | 64.52 | 31.94 | 44.45 | 15.74 | 24.42 | 9.71 | 68.87 | 25.45 | 66.73 | 28.80 |
| 2006 | 37.62 | 17.12 | 27.06 | 12.77 | 64.68 | 30.54 | 44.22 | 14.93 | 25.27 | 10.24 | 69.49 | 25.17 | 67.14 | 27.84 |
| 2007 | 42.33 | 16.47 | 25.80 | 11.90 | 68.13 | 28.37 | 46.67 | 18.74 | 20.18 | 7.10 | 66.85 | 25.84 | 67.47 | 26.80 |
| 2008 | 43.29 | 19.31 | 26.64 | 12.00 | 69.93 | 31.31 | 35.20 | 12.06 | 18.23 | 7.71 | 53.44 | 19.77 | 61.45 | 25.25 |
| 2009 | 37.96 | 13.62 | 25.31 | 9.82 | 63.27 | 23.44 | 41.49 | 14.79 | 20.75 | 8.14 | 62.24 | 22.93 | 62.74 | 23.09 |
| 2010 | 38.35 | 14.60 | 26.91 | 10.79 | 65.26 | 25.39 | 41.12 | 12.99 | 19.61 | 5.75 | 60.73 | 18.75 | 62.92 | 21.92 |
| 2011 | 43.55 | 15.90 | 29.26 | 12.32 | 72.81 | 28.22 | 52.33 | 19.49 | 26.80 | 9.51 | 79.13 | 29.00 | 76.07 | 28.50 |
| 2012 | 63.06 | 24.53 | 32.90 | 13.03 | 95.96 | 37.56 | 34.68 | 11.17 | 28.25 | 11.04 | 62.93 | 22.22 | 78.91 | 29.44 |
| 2013 | 42.26 | 17.71 | 33.25 | 13.21 | 75.51 | 30.92 | 47.33 | 16.84 | 18.16 | 10.44 | 65.49 | 27.28 | 70.34 | 29.08 |
| 2003-2013 | 41.44 | 17.45 | 27.65 | 12.14 | 69.09 | 29.59 | 41.72 | 14.83 | 21.69 | 8.81 | 63.41 | 23.63 | 66.18 | 26.44 |

The age of onset of colon cancer ranged from 17 to 97 years, and the age of onset of rectal cancer ranged from 13 to 99 years; the median values were 74 and 71 years, respectively. The incidence of colon cancer and rectal cancer increased with age, and showed a rapid rise after about 50 years of age; the peak was in the range of 75-80 years of age (Fig.1 CRC incidence rate by age in males and females (2003-2013) (/100 thousand)). Statistically significant gender differences in CRC incidence were found (*P*=0.0431). The standardized incidence rate of colon cancer in males and females was 17.45/10^5^ and 14.83/10^5^, respectively, and that of rectal cancer in males and females was 12.14/10^5^ and 8.81/10^5^, respectively (Table 1 and Fig. 2 Standardized incidence rate by sex (2003-2013) (/100 thousand). Pathological diagnosis accounted for 73.57% of the CRC incidence (Table 2).

**Fig. 1.**
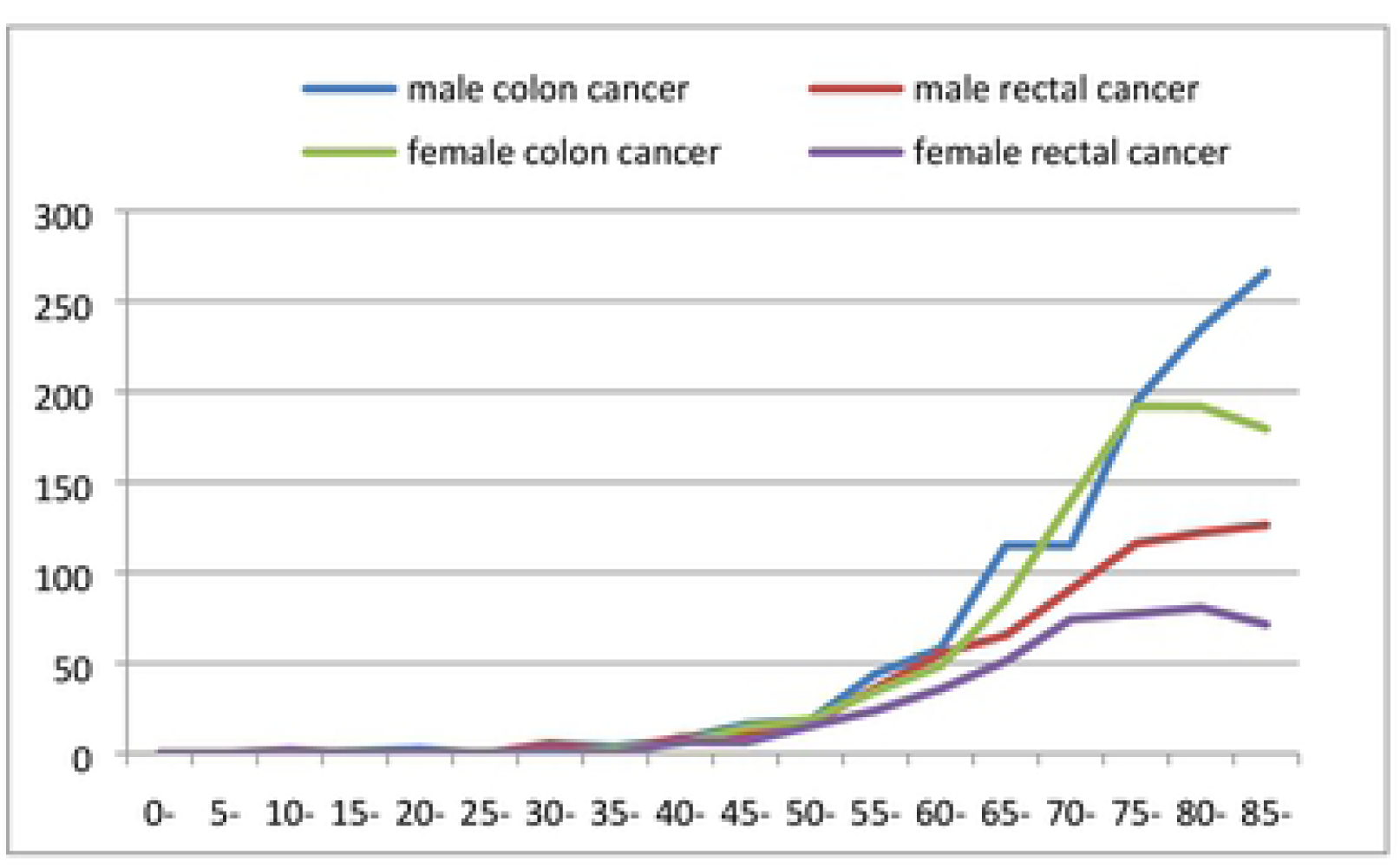
CRC incidence rate by age in males and females (2003-2013) (/100 thousand)

**Table 2.**
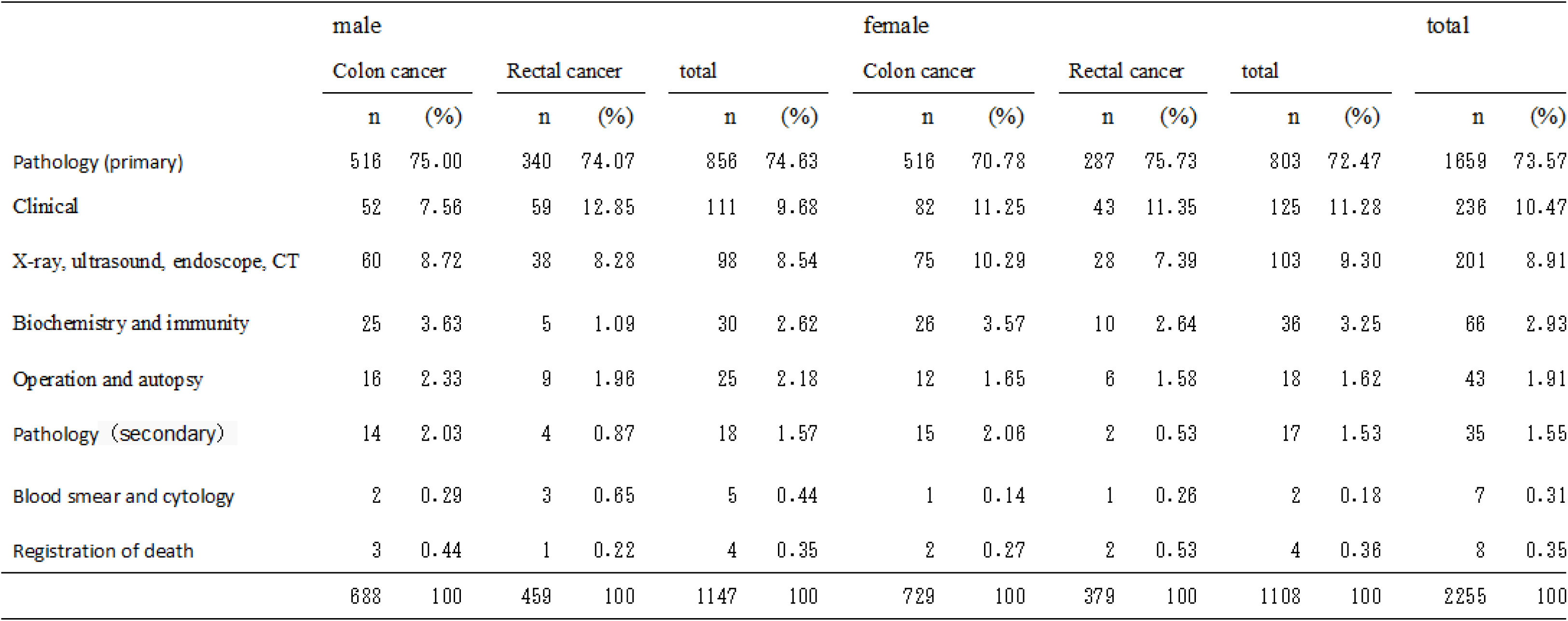
Diagnostic characteristics of highest CRC incidence (2003-2013)

**Fig. 2.**
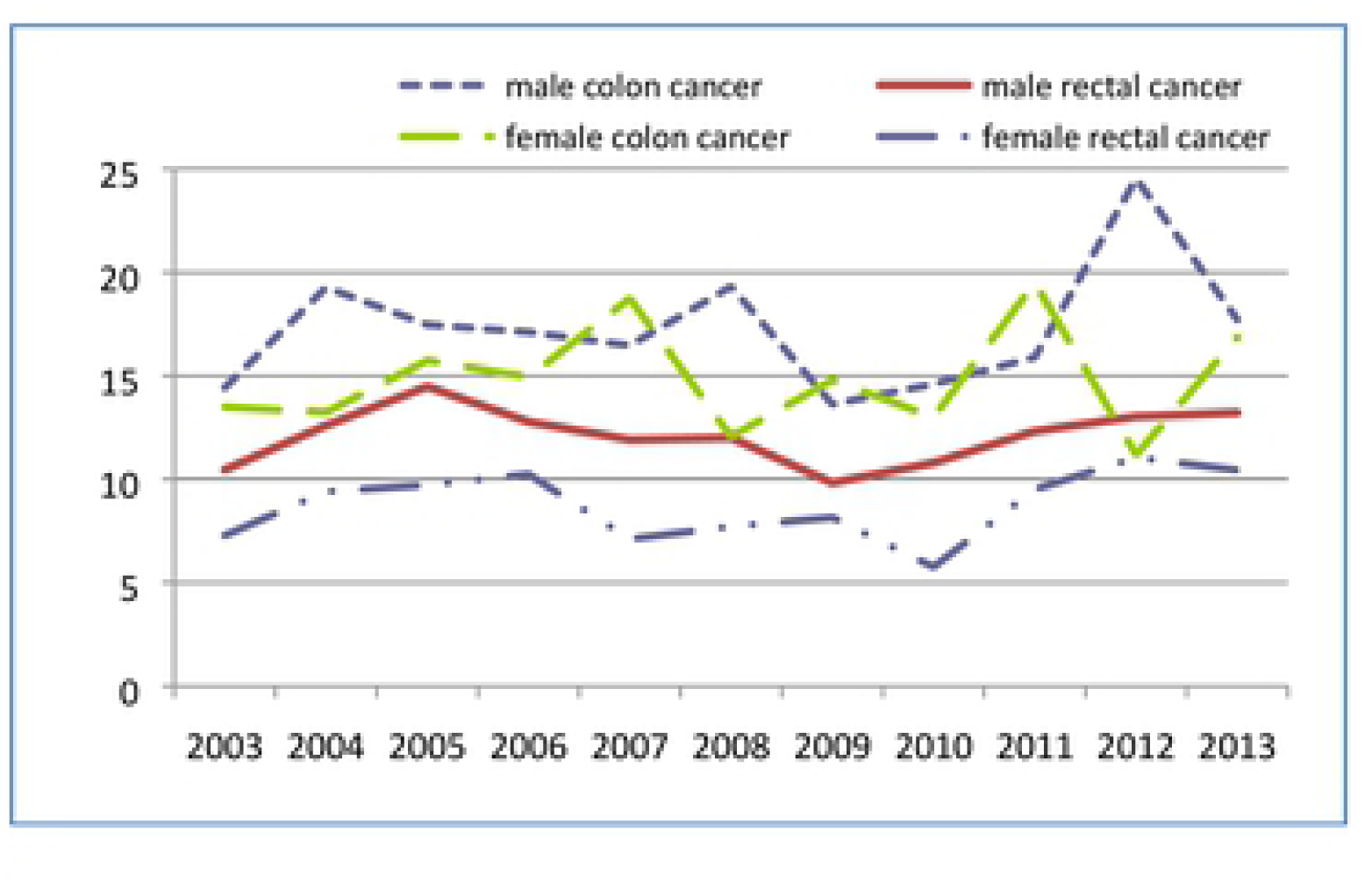
Standardized incidence rate by sex (2003-2013) (/100 thousand)

The crude and standardized CRC mortality rates were 33.10/10^5^ and 10.08/10^5^ (Table 3).

**Table 3.** CRC mortality rate by year and sex (2003-2013) (/100 thousand)

|  | male |  |  |  |  |  | female |  |  |  |  |  | total |  |
| --- | --- | --- | --- | --- | --- | --- | --- | --- | --- | --- | --- | --- | --- | --- |
|  | Colon cancer |  | Rectal cancer |  | total |  | Colon cancer |  | Rectal cancer |  | total |  |  |  |
|  | crude | standard | crude | standard | crude | standard | crude | standard | crude | standard | crude | standard | crude | standard |
| 2003 | 19.91 | 10.15 | 9.95 | 3.55 | 29.86 | 13.71 | 18.09 | 7.31 | 8.44 | 1.64 | 26.53 | 8.95 | 28.17 | 11.00 |
| 2004 | 24.94 | 12.26 | 7.67 | 2.34 | 32.61 | 14.60 | 17.89 | 4.73 | 11.10 | 2.21 | 28.99 | 6.94 | 30.77 | 10.75 |
| 2005 | 24.11 | 10.40 | 12.38 | 3.07 | 36.49 | 13.47 | 18.78 | 5.92 | 8.14 | 1.46 | 26.92 | 7.38 | 31.61 | 10.19 |
| 2006 | 23.76 | 10.57 | 11.22 | 2.97 | 34.98 | 13.54 | 24.64 | 10.01 | 11.37 | 1.95 | 36.01 | 11.96 | 35.50 | 12.60 |
| 2007 | 27.78 | 9.85 | 13.23 | 3.37 | 41.01 | 13.23 | 18.92 | 6.74 | 8.83 | 1.50 | 27.75 | 8.24 | 34.22 | 10.32 |
| 2008 | 17.90 | 8.81 | 12.59 | 3.14 | 30.49 | 11.95 | 13.20 | 5.77 | 13.83 | 2.11 | 27.03 | 7.88 | 28.71 | 10.16 |
| 2009 | 28.26 | 11.16 | 16.82 | 3.95 | 45.07 | 15.11 | 18.34 | 6.61 | 14.55 | 2.19 | 32.89 | 8.80 | 38.48 | 11.61 |
| 2010 | 11.57 | 5.24 | 13.61 | 3.11 | 25.18 | 8.35 | 20.42 | 7.79 | 12.12 | 1.69 | 32.54 | 9.48 | 28.69 | 8.81 |
| 2011 | 28.79 | 10.88 | 12.34 | 2.53 | 41.13 | 13.41 | 17.98 | 6.83 | 17.34 | 2.35 | 35.32 | 9.17 | 37.87 | 11.21 |
| 2012 | 20.78 | 8.30 | 18.70 | 3.83 | 39.49 | 12.13 | 16.86 | 6.17 | 16.86 | 2.27 | 33.72 | 8.44 | 36.14 | 10.13 |
| 2013 | 21.09 | 7.57 | 14.76 | 2.83 | 35.85 | 10.40 | 18.38 | 6.44 | 15.75 | 2.00 | 34.13 | 8.44 | 34.50 | 9.30 |
| 2003-2013 | 22.53 | 9.53 | 12.95 | 2.61 | 35.48 | 12.15 | 18.43 | 6.77 | 12.42 | 1.58 | 30.84 | 8.35 | 33.10 | 10.08 |

The age of death due to colon cancer ranged from 53 to 80 years of age, and the age of death due to rectal cancer ranged from 80 to 100 years of age. The median age of death due to colon and rectal cancers was 73 and 85 years, respectively. Colon cancer mortality increased at about 50 years of age, with peak mortality at about 75 years of age, and then it decreased, exhibiting an “inverted V” shape. Rectal cancer mortality rose rapidly from about 75 years of age, increasing with age (Fig. 3 CRC mortality rate by age in males and females (2003-2013) (/100 thousand)). There were also statistically significant gender differences in CRC mortality (*P*=0.0201). The standardized mortality rate of colon cancer in males and females was 9.53/10^5^ and 6.77/10^5^, respectively, and that for rectal cancer in males and females was 2.61/10^5^ and 1.58/10^5^, respectively (Table 3 and Fig. 4 Standardized mortality rate by sex (2003-2013) (/100 thousand)). Pathological diagnosis accounted for 62.0% of the CRC mortality (Table 4).

**Table 4.**
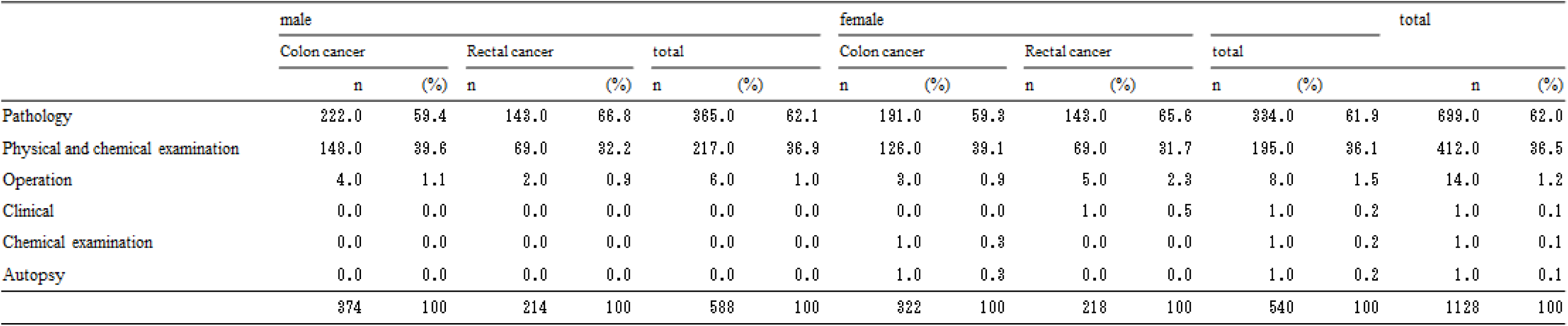
Diagnostic characteristics of highest CRC mortality (2003-2013)

**Fig. 3.**
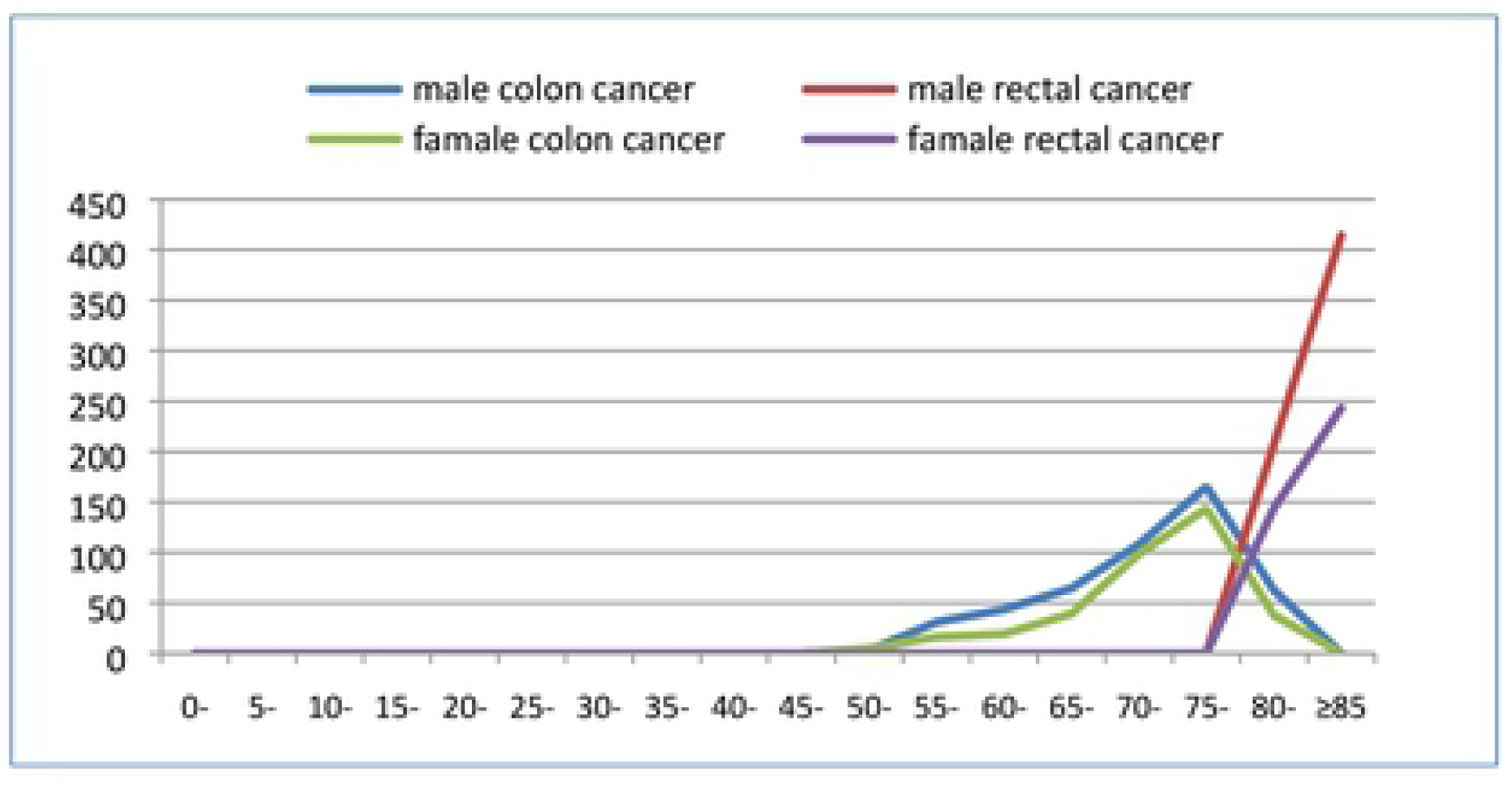
CRC mortality rate by age in males and females (2003-2013) (/100 thousand)

**Fig. 4.**
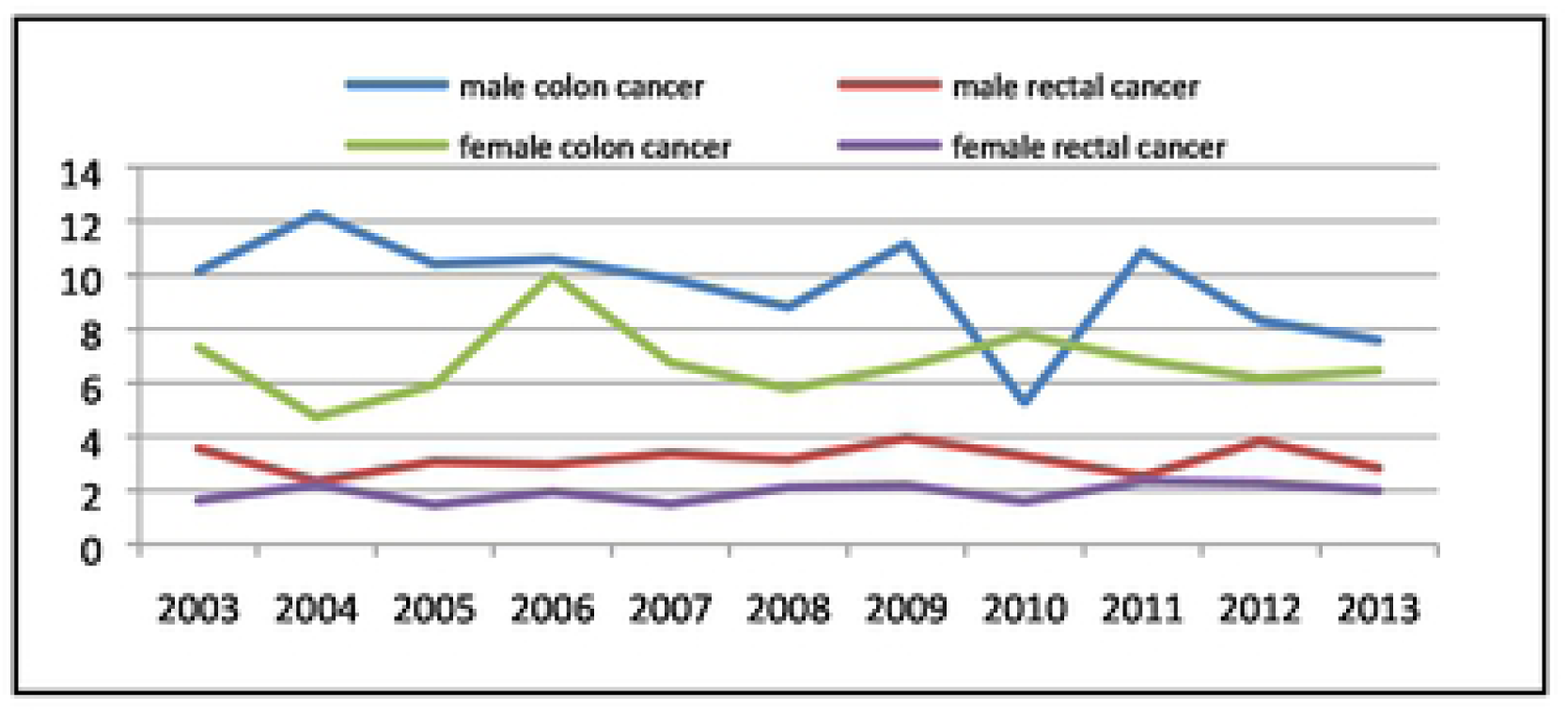
Standardized mortality rate by sex (2003-2013) (/100 thousand)

### Rates of participation in CRC screening in the Jing’an district in 2013

In 2013, for the 82,602 eligible residents, a total of 17,866 underwent CRC screening (21.6%). The compositions of age, sex, marriage and education of the participants are shown in Table 5.

**Table 5.** Characteristics of the 17866 participants.

|  | n | (%) |
| --- | --- | --- |
| Age |  |  |
| 50-59 | 7670 | 42.93 |
| 60-69 | 8367 | 46.83 |
| 70-74 | 1829 | 10.24 |
|  | **17866** | **100** |
| Sex |  |  |
| male | 7052 | 39.47 |
| female | 10814 | 60.53 |
|  | **17866** | **100** |
| Marital status |  |  |
| married | 16175 | 90.54 |
| widowed | 658 | 3.68 |
| divorced | 434 | 2.43 |
| single | 474 | 2.65 |
| unknown | 125 | 0.7 |
|  | **17866** | **100** |
| Education |  |  |
| none | 66 | 0.37 |
| primary school | 685 | 3.83 |
| technical secondary school/middle school | 12802 | 71.66 |
| undergraduate/postsecondary | 4211 | 23.57 |
| graduate | 102 | 0.57 |
|  | **17866** | **100** |

The total positive rate by either the HRFQ and/or the FIT was 16.28% (2909/17,866). The HRFQ positive rate was 7.49% (1339/17,866), while the FIT positive rate was 9.9% (1761/17,866) (Table 6).

**Table 6.**
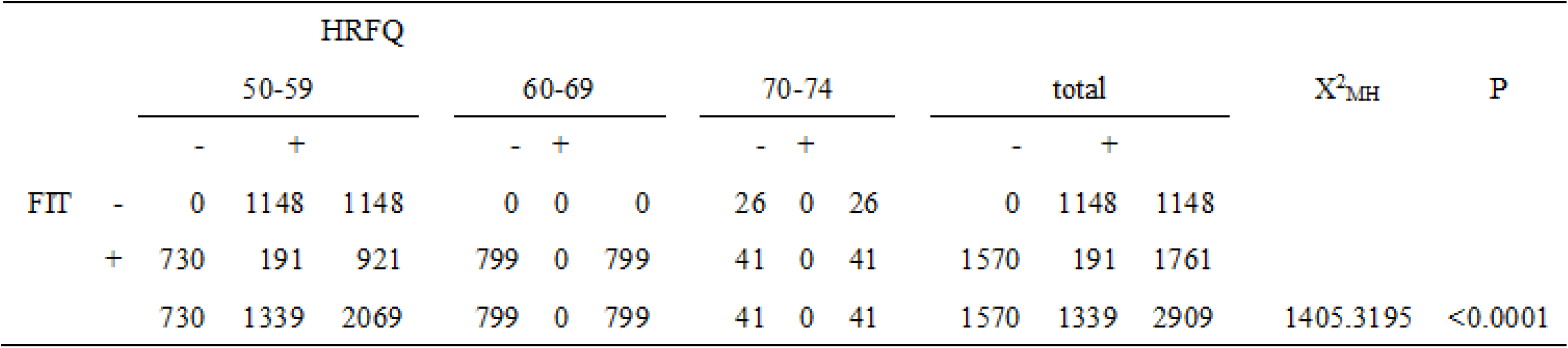
Positive screening questionnaire and FIT.

The compositions of age, sex, marriage and education of those who screened positive are shown in Table 7.

**Table 7.** Characteristics of the 2909 subjects who screened positive.

|  | group | n | (%) |
| --- | --- | --- | --- |
| Age |  |  |  |
|  | 50-59 | 2069 | 71.1 |
|  | 60-69 | 799 | 27.5 |
|  | 70-74 | 41 | 1.4 |
| Sex |  |  |  |
|  | male | 966 | 33.2 |
|  | female | 1943 | 66.8 |
| Marital status |  |  |  |
|  | married | 2591 | 89.1 |
|  | divorced | 81 | 2.8 |
|  | widowed | 148 | 5.1 |
|  | single | 71 | 2.4 |
|  | unknown | 18 | 0.6 |
| Education |  |  |  |
|  | none | 10 | 0.3 |
|  | primary school | 101 | 3.5 |
|  | technical secondary school/middle school | 2022 | 69.5 |
|  | undergraduate/postsecondary | 760 | 26.1 |
|  | graduate | 16 | 0.6 |

### Low rates of participation in follow-up colonoscopy in the Jing’an district in 2013

The 2909 individuals who screened positive for CRC were referred to four designated hospitals for follow-up colonoscopy (CSPY). However, only 508 (17.5%) underwent CSPY in 2013 (Fig. 5 Flow-chart of the screening program). The outcome of CSPY was normal for 282 (55.5%), abnormal for 210 (41.3%), and unknown for 16 (3.1%). Eight cases were diagnosed as CRC (1.6%), and 142 cases were diagnosed as precancerous lesions (28.0%) (Table 8).

**Table 8.**
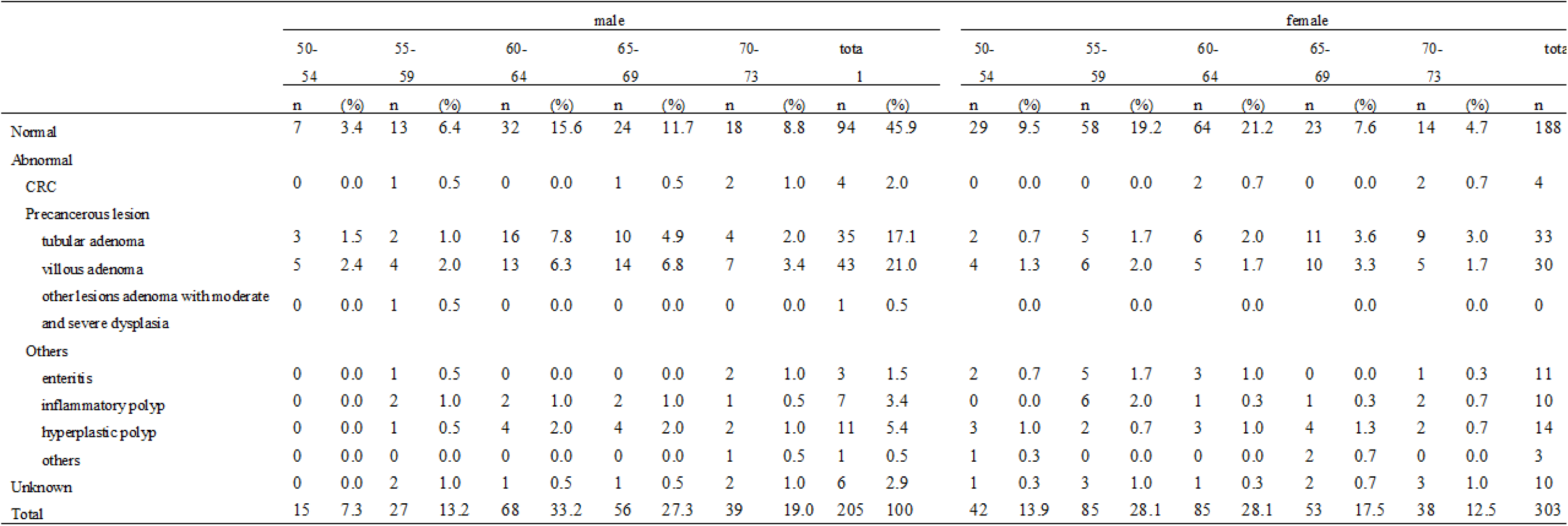
Clinical characteristics of the 508 colonoscopy cases.

**Fig. 5.**
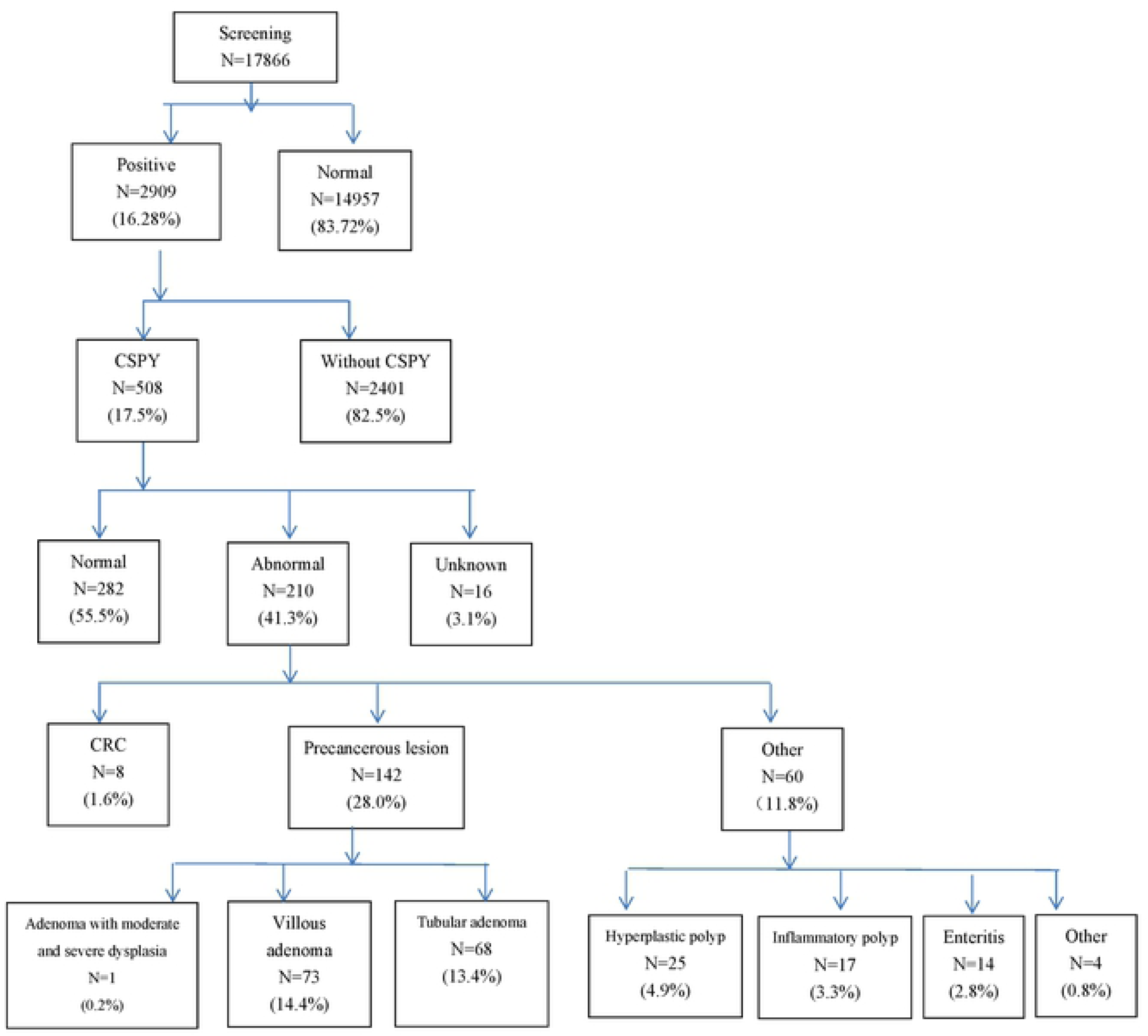
Flow-chart of the screening program.

The diagnostic stages of CRC are shown in Table 9.

**Table 9.** TNM classification of the 8 CRC cases.

| n | T | N | M |
| --- | --- | --- | --- |
| 1 | T<sub>1</sub> | N<sub>0</sub> | M<sub>0</sub> |
| 2 | T<sub>3</sub> | N<sub>0</sub> | M<sub>0</sub> |
| 2 | T<sub>4a</sub> | N<sub>0</sub> | M<sub>0</sub> |
| 1 | T<sub>4a</sub> | N<sub>1</sub> | M<sub>0</sub> |
| 2 | Unknown |  |  |

## Discussion

### CRC incidence and mortality in the Jing’an district were similar to those of the entire Shanghai area, which is among the areas of high CRC incidence and mortality in China

The standardized CRC incidence and mortality rates assessed between 2003 and 2013 were 26.44/10^5^ and 10.08/10^5^, respectively in the Jing’an district, similar to those between 2003 and 2007 (22.23/10^5^ and 12.96/10^5^) and (20.19/10^5^ and 9.22/10^5^) in the entire Shanghai area calculated by the municipals. From other surveys of the data calculated by the municipals in other cities and counties of China between 1998 and 2002, the CRC standardized incidence rates in Shanghai males and females were 20.4/10^5^ for males and 25.5/10^5^ for females, and the CRC standardized mortality rates were 15.1/10^5^ for males and 11.6/10^5^ for females. These rates of CRC incidence and mortality in Shanghai were higher than those in the other core cities in China. Combined with the results of the present study, it is evident that Shanghai is a city of high CRC incidence and mortality among Chinese core cities, and the Jing’an district appears to be representative of an urban district of Shanghai. Even though the rates of CRC incidence and mortality in the Jing’an district were relatively high compared with other Chinese areas, they are still lower than those in the Western countries.[1-3] In the US, the CRC incidence rate in 2010 was estimated at 40/10^5^, and the CRC mortality rate was estimated at 15/10^5^.[18] However, in China, other data have suggested gradual increases in CRC incidence and mortality, presumably owing to a shift to a more Western lifestyle with regard to food, physical activity, etc. These tendencies in age-related increases and gender differences in CRC incidence and mortality in the Jing’an district [Table 1-2, Fig. 1-4] were consistent with those in past reports in other areas in China and other countries.[1,19] The gender difference in China might be due, in part, to a greater awareness of prevention in females than in males.

### Combined use of FIT and HRFQ was effective in CRC screening programs as an initial screening method

For CRC screening, we utilized the Consensus in Early Diagnosis and Early Treatment, Comprehensive Prevention of China CRC Screening approach, using HRFQ, and FIT was used instead of guaiac fecal occult blood test (gFOBT).[20,21]. The FIT test was an immunoassay to detect faecal human haemoglobin at very low concentrations, and was considered more sensitive than the gFOBT.[22] Moreover, FIT is advantageous in providing a quantitative result that enables users to set acceptable rates of screening positives.[23] Though further improvement in the FIT assessment and development of other assessments are in progress[24-29], the FIT used in this study appears reliable and appropriate for CRC screening at this stage.

### Follow-up colonoscopy (CSPY) is indispensable for reducing CRC mortality in individuals who screen positive for CRC

CSPY is considered the gold standard for the diagnosis of CRC and polyps to prevent death, and it is the most effective strategy to identify polyps at the initial stages. Moreover, CSPY is a less invasive method to remove colorectal adenomas immediately upon identification. Thus, it is evident that CSPY surveillance is beneficial for most patients with intermediate-risk adenomas.[30] At present, other screening methods include ultrasound endoscopy (EUS), positron-emission tomography (PET) and genetic diagnosis.[31]. More recently, colon capsule endoscopy (CCE) was proposed for CRC screening by the European Digestive Disease Week and the World Congress of Gastrointestinal Disease.[32]. However, with CCE, it is impossible to obtain histological samples, and it is more expensive, although it is much more acceptable than CSPY in terms of pain expectations.[33]. In this study, among the 508 patients who underwent follow-up CSPY, 210 (41.3%) were judged as abnormal, and 8 (3.1%) were diagnosed with CRC (Table 7). Given that previous data indicated that only 62.0% of CRC deaths had a pathological diagnosis, follow-up CSPY is quite effective for the prevention of CRC mortality.

### Low participation rate in follow-up CSPY after CRC screening

Regardless of the significance of follow-up CSPY to reduce CRC mortality, the rate of participation in follow-up CSPY after CRC screening in 2013 was only 17.5% (508/2909) in this study. From the registry of the Shanghai CRC Screening Information System, the rates of participation in follow-up CSPY from 2014 to 2016 in the Jing’an district were also only 11.65% (72/618) in 2014 and 5.43% (59/1086) in 2015. Such annual decreases in the rates of participation were also recorded for the initial CRC screening [18.28% (618/3380) in 2014 and 26.92% (1086/4034) in 2015]. Moreover, a similarly low rate of follow-up CSPY participation was found between 2012 and 2013, as only 9.51% (1000/10512) was reported in the Xuhui District.[34] These low rates of follow-up CSPY screening after positive results of CRC screening in China, at least in the urban areas, are striking, since much better rates of participation in follow-up CSPY have been reported in the US (more than 50%) [35] and Japan (69.1% = 10593/153320 positives, among 2535814 participants in CRC screening in 2016).[36]

Similarly, in a study of the national screening programme in England, about 60% of the individuals who screened positive for CRC participated in follow-up CSPY screening in two years later.[37] The exact reason for such a low rate of participation in follow-up CSPY in China remains to be studied, but several issues could be responsible. CSPY requires several steps, such as bowel preparation and a complex procedure with possible complications, which may deter individuals who screen positive for CRC from undergoing CSPY. It is also suggested that even if the initial CRC test is abnormal, some individuals may feel that further follow-up with CSPY screening is unnecessary.

### CRC and follow-up CSPY screenings are effective in reducing CRC incidence and preventing CRC mortality

Decreasing CRC mortality rates have been observed in a large number of countries worldwide. This trend is assumed to be attributed mainly to CRC screening. [38,39]. For example, CRC mortality rates declined by 47% among men and by 44% among women from 1990 to 2015 in the United States, and in 1995, the US Preventive Services Task Force (USPSTF) endorsed screening with faecal occult blood testing (FOBT) and flexible sigmoidoscopy. New guidelines released by the USPSTF in 2016 emphasized the benefits of CRC screening in average-risk individuals, and listed 8 possible screening programs that could reduce CRC mortality.[40] In Germany, within 10 years of the introduction of screening CSPY, the incidence of bowel cancer in persons over the age of 55 fell by 17–26%, after having risen steadily over the preceding decades.[41] The UK bowel cancer awareness campaigns appeared to have improved public awareness of CRC and encouraged symptomatic individuals to seek urgent medical attention.[42] In Australia, Australian National Bowel Cancer Screening Program (NBCSP) invitees had less risk of dying from CRC[43], and were more likely to have less-advanced disease when diagnosed compared with noninvitees, and the potential benefits of a program similar to the NBCSP may become available to an increasing number of countries.

The present study indicate that efforts are required for rigorous improvement in CRC and follow-up screenings in China. Regarding the low rates of CRC screening, some factors affecting participation like socio-demographic characteristics (educational level, health insurance, and knowledge of CRC and CRC screening), psychological factors (perceived severity of CRC, susceptibility to CRC, and barriers to screening), and contact with medical provider (physician recommendation) could be considered.[44] To improve participation in CRC screening in China, more studies are clearly needed to address these points including a longer follow-up of the present study, to better inform the citizens regarding the benefits of CRC screening.

## Conclusion

The incidence and mortality of CRC were high in the Jingan district, a representative urban Chinese area. Nevertheless, the rates of participation in CRC screening and in follow-up CSPY for individuals who screened positive were quite low when compared to those reported in other countries. Therefore, improvement in participation in CRC screening is important to significantly lower the incidence and mortality of CRC.

## Acknowledgments

This study was funded by the Jingan District Shanghai Health and Family Planning Commission Programs, number JWXK201209. The study also received financial support from the Jingan District Government, and from Tokiwa Foundation for SK. We would like to thank all of the related staff in the Fudan University School of Public Health, Shanghai, China. We also thank the members of Medical Governance Research Institute, Tokyo, Japan and Ms. Rongrong Liang for supports, as well as personal assists.

